# Starvation Decreases Immunity and Immune Regulatory Factor NF-κB in the Starlet Sea Anemone *Nematostella vectensis*

**DOI:** 10.1101/2022.06.09.495518

**Authors:** Pablo J. Aguirre Carrión, Niharika Desai, Joseph J. Brennan, James E. Fifer, Sarah W. Davies, Thomas D. Gilmore

## Abstract

Lack of proper nutrition (malnutrition) or the complete absence of all food (starvation) have important consequences on the physiology of all organisms. In many cases, nutritional status affects immunity, but, for the most part, the relationship between nutrition and immunity has been limited to studies in vertebrates and terrestrial invertebrates. Herein, we describe a positive correlation between nutrition and immunity in the sea anemone *Nematostella vectensis*. Gene expression profiling of adult fed and starved anemones showed downregulation of many genes involved in nutrient metabolism and cellular respiration, as well as immune-related genes, in starved animals. Starved adult anemones also had reduced protein levels and DNA-binding activity of immunity-related transcription factor NF-κB. Starved juvenile anemones had increased sensitivity to bacterial infection and also had lower NF-κB protein levels, as compared to fed controls. Weighted Gene Correlation Network Analysis (WGCNA) revealed significantly correlated gene networks that were inversely associated with starvation. Based on the WGCNA and a reporter gene assay, we identified *TRAF3* as a likely NF-κB target gene in *N. vectensis*. Overall, these experiments demonstrate a correlation between nutrition and immunity in a basal marine metazoan, and the results have implications for the survival of marine organisms as they encounter changing environments.

**Significance Statement:** Adequate nutrition is required to sustain proper biological function. One factor threatening many marine organisms, as a result of modern day anthropogenic environmental changes, is nutrient availability. Here, we characterize transcriptional changes following food deprivation in the cnidarian model sea anemone *Nematostella vectensis*. We show that starvation is correlated with decreased expression of genes associated with nutrient metabolism and immunity, among others. Moreover, starvation reduces the level of expression and activity of immune regulatory transcription factor NF-κB and causes anemones to have increased susceptibility to bacterial infection. These results demonstrate that this basal organism responds at the transcriptional level to the absence of food, and that, in addition to changes in metabolic factors, starvation leads to a reduction in immunity.

## Introduction

Maintenance of caloric needs is a requirement across the tree of life. Food scarcity is a challenge most heterotrophs encounter at various points during their lives – a situation that contributes to natural selection. The lack of proper nutrition (malnutrition) or the complete absence of food (starvation) has important consequences on an organism’s physiology. Starvation can lead to the slowing of metabolism^1^, and nutrition is one of the many factors that determine immune status of organisms^2^. Indeed, the adverse impact of poor nutrition on the immune system, including its inflammatory component, is well documented in vertebrates and some terrestrial invertebrates^3-11^. However, the interplay between nutrition and immunity has, for the most part, only been described in more derived organisms from insects to mammals.

The cnidarian model starlet sea anemone *Nematostella vectensis* (*Nv*) shows remarkable adaptability – being able to survive across a wide range of salinity, pH, and temperature – and it can be readily maintained and studied in the laboratory^12^. *Nv* also has exceptional regenerative abilities, being able to replace its entire oral end in ∼6 days following bisection^13,14^. Importantly, *Nv* can withstand periods of starvation for over a month, where its body size decreases in response to the lack of caloric intake while maintaining body proportionality^15^, suggesting that *Nv* can physiologically respond to changes in food availability. This ability is not limited to adult *Nv*, as previous work has shown that the timing of tentacle development in young *Nv* polyps is intertwined with feeding^16^. *Nv* is also distantly related to reef-building corals, where food availability has been shown to mitigate negative consequences of climate change^17-19^.

Proper resource allocation is vital for survival. Among energetic needs, the immune system is metabolically costly and subject to regulation^20^. Transcription factor NF-κB (nuclear factor kappa B) has been intensively studied in animals from insects to vertebrates for its involvement in development and immunity^21^. Although the basic NF-κB pathway in *Nv* has been characterized^22^ and has been linked to development^23^ and immunity^24,25^, the relationship between nutrition and immunity in *Nv* – or any cnidarian – remains vastly unexplored. Mammals have five different NF-κB proteins and flies have three, all of which are involved in some aspect of immunity^21^. Most basal organisms, including sponges and cnidarians, have single NF-κB proteins; however, the role of NF-κB in these organisms is less clear. Although NF-κB is regulated by post-translational mechanisms in organisms more derived than flies, there is evidence that NF-κB activity is regulated at the transcriptional level in many basal metazoans^26-28^. Examples of NF-κB target genes include cytokines and other immune response factors in mammals, and anti-microbial proteins such as cecropins in insects^29-30^. Furthermore, NF-κB signaling has been shown to play a key role in metabolic disease given its role in inflammation and the association of chronic inflammation with diseases such as obesity and diabetes^6^.

Herein, we have investigated the relationship between nutrition and immunity in *Nv* using gene expression profiling, characterization of transcription factor NF-κB, and susceptibility to infection with a bacterial pathogen. These results directly connect immunity and nutrition in a cnidarian, suggesting that nutritional status and immunity are evolutionarily linked processes.

## Results

### Genome-wide transcriptomic changes in *N. vectensis* in response to starvation

To assess the impact of reduced nutrition on *N. vectensis* (*Nv*), we performed genome-wide gene expression profiling using TaqSeq on eight clonal anemone pairs wherein single animals from clonal pairs were either fed routinely or starved for four weeks prior to RNA isolation (Fig. 1a). Clonal pairs were used because preliminary experiments showed heterogeneity in protein expression when comparing anemones of differing genetic backgrounds. To avoid positional effects after regeneration, we included animals regenerated from aboral and oral ends in both fed and starved groups (Extended Data Table 1). Raw sequencing reads from all 16 anemones ranged from 5.2 to 9.2 million. Alignment values for all samples ranged from 74.2 to 77.4%. *DESeq2* identified 711 significant differentially expressed genes (DEGs) between starved and fed anemones while controlling for genetic background (*FDR adjusted p-value* < 0.1): 118 genes were upregulated and 593 genes were downregulated in starved relative to fed anemones (Fig. 1b). A full list of the differentially expressed genes is presented in Extended Data Dataset 1.

**Figure 1.**
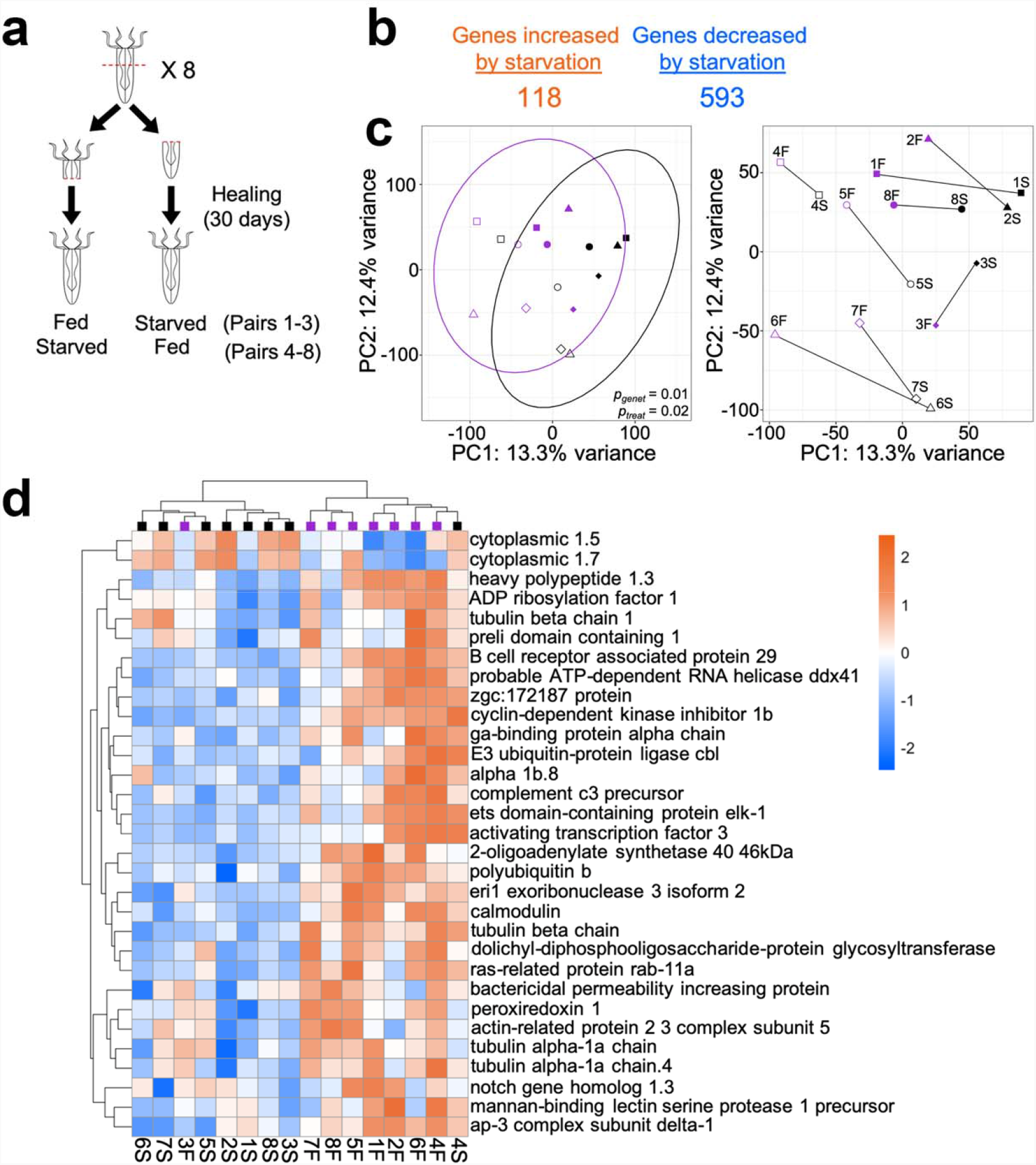
Changes in mRNA expression induced by starvation of adult *Nematostella*. **a**, Schematic of experimental design. Eight adult anemones were bisected perpendicular to their body column and allowed to heal for 30 days before subjecting them to starvation. After two weeks of feeding or starvation, RNA was isolated and subjected to TaqSeq analysis. **b**, Number of significantly (*FDR adjusted p-value* < 0.1) increased and decreased DEGs after 30 days of starvation. **c**, Principal component analysis (PCA) of *rlog* normalized gene expression profiles in *Nematostella*. Color denotes feeding status, i.e., either fed (purple) or starved for 14 days (black). Samples with the same symbol are clonal pairs. PERMANOVA results are shown for genetic background (*p*_*genet*_) and feeding treatment (*p*_*treat*_). PCA plot with ellipses (left) denoting multivariate t-distribution for both feeding status groups. The right shows the same plot with lines connecting samples that are clonal pairs. **d**, Hierarchical clustered heatmap of DEGs annotated with the “Immune Response” (*GO:0006955*) GO term and a *p-value* < 0.01. Colors indicate direction (blue, decreased with starvation; orange, increased with feeding) and magnitude of response through difference in expression relative to mean expression across samples. Colored squares at the top indicate feeding regime (purple, fed; black starved).

To assess overall transcriptional differences between fed and starved anemones, *rlog-* normalized gene expression data were used for Principal Components Analysis (PCA), which showed that nutritional status had a significant influence on gene expression (Fig. 1c, left; *Adonis* PERMANOVA *p*_*treat*_ = 0.02). Additionally, the effect of genet (i.e., genetic background) also had a significant impact on gene expression (Fig. 1c, *Adonis* PERMANOVA *p*_*genet*_ = 0.01). These differences in gene expression between fed and starved animals were also seen when comparing individual clonal pairs where each starved anemone showed a similar rightward shift along PC1 compared to its fed counterpart (Fig. 1c, right).

### GO pathway response to starvation

To explore the functional response of *Nv* to starvation, a Gene Ontology (GO) enrichment analysis of “Biological Process” terms associated with significant DEGs (Extended Data Fig. 1) was performed using ranked *p*-values. Among genes showing positive log-fold changes following starvation, three prominent Biological Process trends emerged: RNA processing (i.e., *RNA processing GO:0006396, RNA splicing GO:0008380, mRNA metabolic process GO:0016071*), DNA processing (i.e., *DNA metabolic process GO:0006259, DNA replication GO:0006260, DNA repair GO:0006281*), and chromosome organization (i.e., *chromatin remodeling GO:0006338, chromatin organization GO:0006325, covalent chromatin modification GO:0016569*).

Consistent with results from the DEG analysis (Fig. 1b), more GO terms were underrepresented in starved anemones. Many of the underrepresented GO terms reflected the nutrient-deprived status of these animals, with terms related to nutrient metabolism (i.e., *response to nutrient GO:0007584, lipid metabolic process GO:0006629, carbohydrate metabolic process GO:0005975, cellular amino acid metabolic process GO:0006520*) and cellular respiration (i.e., *electron transport chain GO:0022900, proton transmembrane transport GO:1902600*) (Extended Data Fig. 1). Thus, these downregulated GO terms indicate that the anemones were indeed starved.

Of note, many immune-related terms were underrepresented following starvation (i.e., *immune response GO:0006955, defense response GO:0006952, antigen processing and presentation GO:0019882*), as well as some oxidative stress-related terms (i.e., *reactive oxygen species metabolic process GO:0072593, hydrogen peroxide metabolic process GO:0042743*) (Extended Data Fig. 1). Figure 1d presents expression data for the 31 significant (*FDR adjusted p-value* < 0.1) annotated DEGs associated with the *immune response* GO term. This group included *Nv* homologs of Notch and Elk1, and members of the complement system. Samples clustered according to feeding regime, with the exception of the fed clone 3 and starved clone 4 (anemones 3F and 4S).

### Starvation increases the susceptibility of *N. vectensis* polyps to bacterial infection

Given that GO terms associated with immunity were underrepresented in starved anemones, we hypothesized that *Nv*’s ability to withstand bacterial challenge would be reduced in starved anemones. *Pseudomonas aeruginosa* is a Gram-negative bacterium that is pathogenic in a variety of hosts including some plants, invertebrates, and vertebrates^31,32^. Recently, *P. aeruginosa* strain PA14 was also shown to be pathogenic for juvenile *Nv*^25^. To determine if fed and starved anemones have different susceptibility to *P. aeruginosa*, we infected a total of 60 fed and 60 starved 40-day old juvenile anemones over three separate experiments with an average of ∼4.5 × 10^8^ CFU/ml of PA14. We then visually monitored disease progression daily based on tissue degradation and death. We found that starved anemones died at a significantly faster rate than their fed counterparts (Kaplan-Meier *p* > 0.01, N = 24, Fig. 2), and that this pattern was reproducible across trials (Extended Data Fig. 2 a & b for two additional trials). Therefore, consistent with reduced expression of immunity-related genes in adults, starved juvenile *Nv* have increased susceptibility to the effects of a bacterial pathogen.

**Figure 2.**
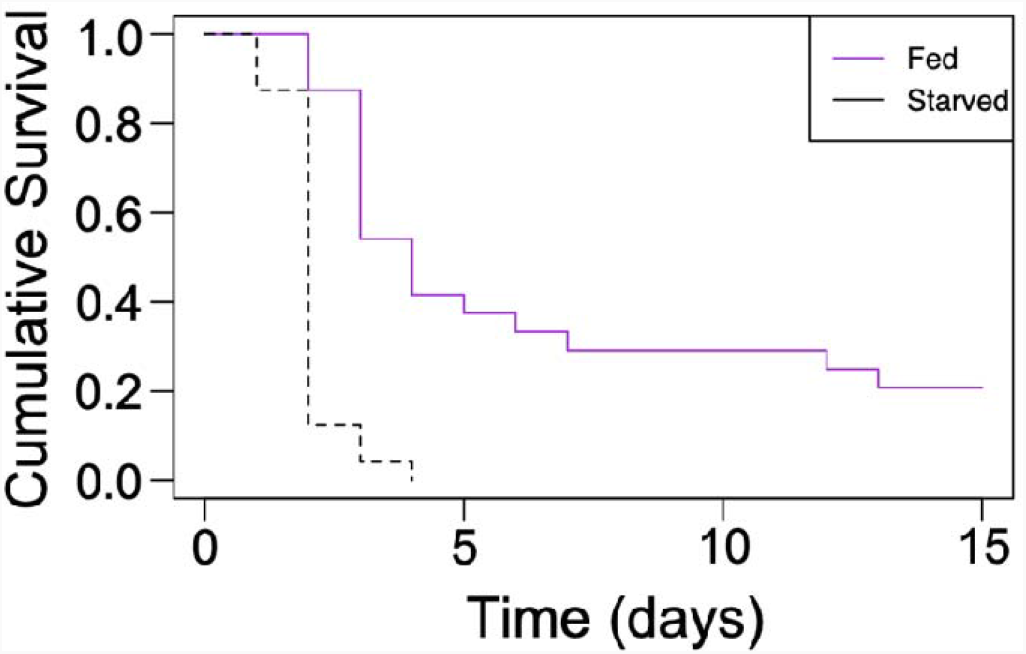
Starved anemones have increased susceptibility to *Pseudomonas aeruginosa* infection-induced death. Two-week old anemones were either fed on a regular schedule (purple) for 30 days or starved (black), and were then infected with 6.8 × 10^8^ CFU/ml of *P. aeruginosa* at 28°C. Survival was monitored daily for 15 days and recorded. N = 24 for both feeding regimes. Significance was determined using Kaplan-Meier statistics.

### NF-κB protein and DNA-binding activity are reduced in starved anemones

Transcription factor NF-κB has a broad role in immunity across Metazoa^21,33,34^. We have also previously shown that NF-κB protein is expressed in juvenile and adult *Nv*, with expression starting as early as 30 h post-fertilization^22,35^. Given the upregulation of NF-κB following infection with *P. aeruginosa*^25^ and that several immunity-related genes were downregulated in our starved anemone gene expression data, we aimed to determine whether starvation had an effect on NF-κB signaling. We first hypothesized that NF-κB transcripts would be downregulated under starvation in *Nv*, and while gene expression patterns in fed and starved adult *Nv* showed that NF-κB transcripts were downregulated in starved anemones (log_2_Fold Change = -0.96, *raw p-value* = 0.01), this trend was not significant after FDR correction.

We next compared NF-κB protein and DNA-binding levels in starved *vs* fed adult anemones. To do this, we again generated clonal pairs (N = 3) of animals (by bisection and regeneration), which were then fed or starved for two weeks. To compare NF-κB protein levels, Western blotting was performed on whole animal extracts from fed and starved clonal pairs. On average, starved anemones had ∼70% less NF-κB protein than their fed counterparts (Fig. 3a). To compare NF-κB DNA-binding activity, an electromobility shift assay (EMSA) was performed using extracts from three fed *vs* starved anemone pairs and a ^32^P-labeled κB-site probe that we have previously shown can be specifically bound by Nv-NF-κB^36^. Consistent with the decrease in protein levels, starved anemones had less NF-κB DNA-binding activity than their fed counterparts (Fig. 3b).

**Figure 3.**
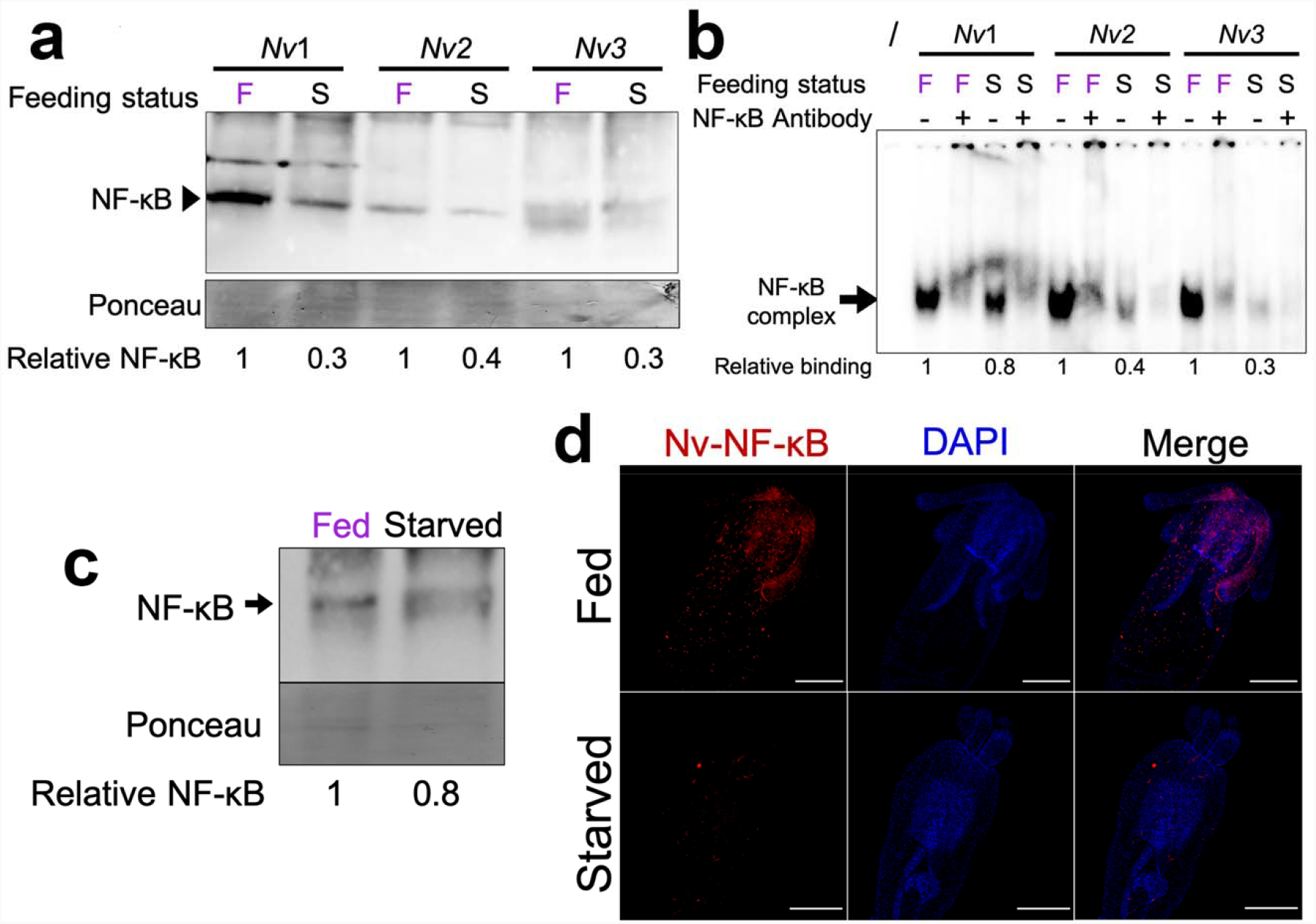
Starvation causes decreased levels of NF-κB. **a**, Two-week fed or starved clonal *Nv* pairs were lysed and 100 µg of protein was electrophoresed on a 7.5% SDS-polyacrylamide gel and subjected to Western blotting with anti-Nv-NF-κB antiserum. Of note, *Nv*’s NF-κB is present as two naturally occurring alleles, one of which migrates slightly lower (36) and can be observed as a separate band in clone *Nv3*. NF-κB protein levels were normalized to Ponceau staining and quantified using ImageJ, and the relative NF-κB protein levels are indicated for each anemone pair. Both naturally occurring alleles of Nv-NF-κB were accounted for during quantification. **b**, Starvation decreases Nv-NF-κB DNA-binding activity. 100 µg of protein extract was incubated with a ^32^P-labelled κB probe (GGGAATTCCC) and samples were analyzed by EMSA. Feeding status is labeled as follows: F, fed and S, starved. Where indicated, samples were also incubated with anti-NF-κB antiserum for super shifting. First lane denoted by ‘/’ is the probe with no protein extract. **c**, Western blotting of juvenile anemones. Forty-day old *Nv* that had been either fed ground *Artemia* or never fed, were pooled together (100 individuals for each feeding regimen), lysed directly in SDS buffer, and lysates were subjected to Western blotting. Protein levels were normalized to Ponceau staining and quantified using ImageJ, and NF-κB protein level is relative to that seen in fed animals. Both naturally occurring alleles of Nv-NF-κB were accounted for during quantification. **d**, Whole-mount immunofluorescence was performed on 40-day old polyps that were either fed for 30 days or never fed. Nv-NF-κB was detected with a custom antibody and visualized with Texas Red-labeled secondary antibody (red, left panels) and nuclei with DAPI (blue, middle panels). Scale bars, 100 μm. All images were taken on a Nikon C2 Si. Images are representatives of both feeding conditions (N = 6 for both treatments; Extended Data Table 2). Statistical significance was determined with an unpaired *t* test (*p* = 0.04).

Because of the increased susceptibility of starved juvenile anemones to bacterial infection (Fig. 2), we were interested in determining whether the overall decrease in NF-κB protein during starvation was also observed in juvenile *Nv*. To do this, we performed anti-Nv-NF-κB Western blotting and immunohistochemistry on 40-day old *Nv* that had been fed regularly or never fed. To analyze Nv-NF-κB protein levels in juvenile *Nv*, we pooled 100 fed or starved polyps, lysed each pool by boiling in SDS sample buffer, and analyzed those lysates by Western blotting. Results showed that starved polyps had ∼20% less NF-κB protein than fed anemones (Fig. 3c). In addition, starved juveniles had ∼60% fewer detectable NF-κB-positive cells by immunohistochemistry (Unpaired *t-*test *p* < 0.05) (Fig. 3d; Extended Data Table 2). Taken together, these results indicate that starvation causes the downregulation of Nv-NF-κB in adult anemones, which results in decreased NF-κB protein and DNA-binding activity, and that decreased NF-κB protein expression is also seen in juvenile anemones, which have increased susceptibility to bacterial infection.

### Gene co-expression network analysis reveals possible cnidarian immune gene network

NF-κB is known for its role as master regulator of immunity^33,34^; this role can include the regulation of other members in the NF-κB signaling pathway. One such member, TNF receptor-associated factor 3 (*TRAF3), h*as been previously observed to be upregulated with NF-κB in *Nv*^25^. Indeed, we found the *Nv TRAF3* gene has three predicted strong NF-κB sites within 1000 bp of its transcription start site (TSS) (Extended Data Fig. S5). All three of these predicted NF-κB sites can be bound by bacterially expressed Nv-NF-κB in an electrophoretic mobility shift assay (Fig. 4a). Furthermore, a luciferase reporter plasmid containing the upstream promoter region of *Nv*-*TRAF3* can be activated by Nv-NF-κB when co-transfected in HEK 293 cells (Fig. 4b), and mutation of the three NF-κB binding site abolished its ability to be activated by Nv-NF-κB (Fig. 4b).

**Figure 4.**
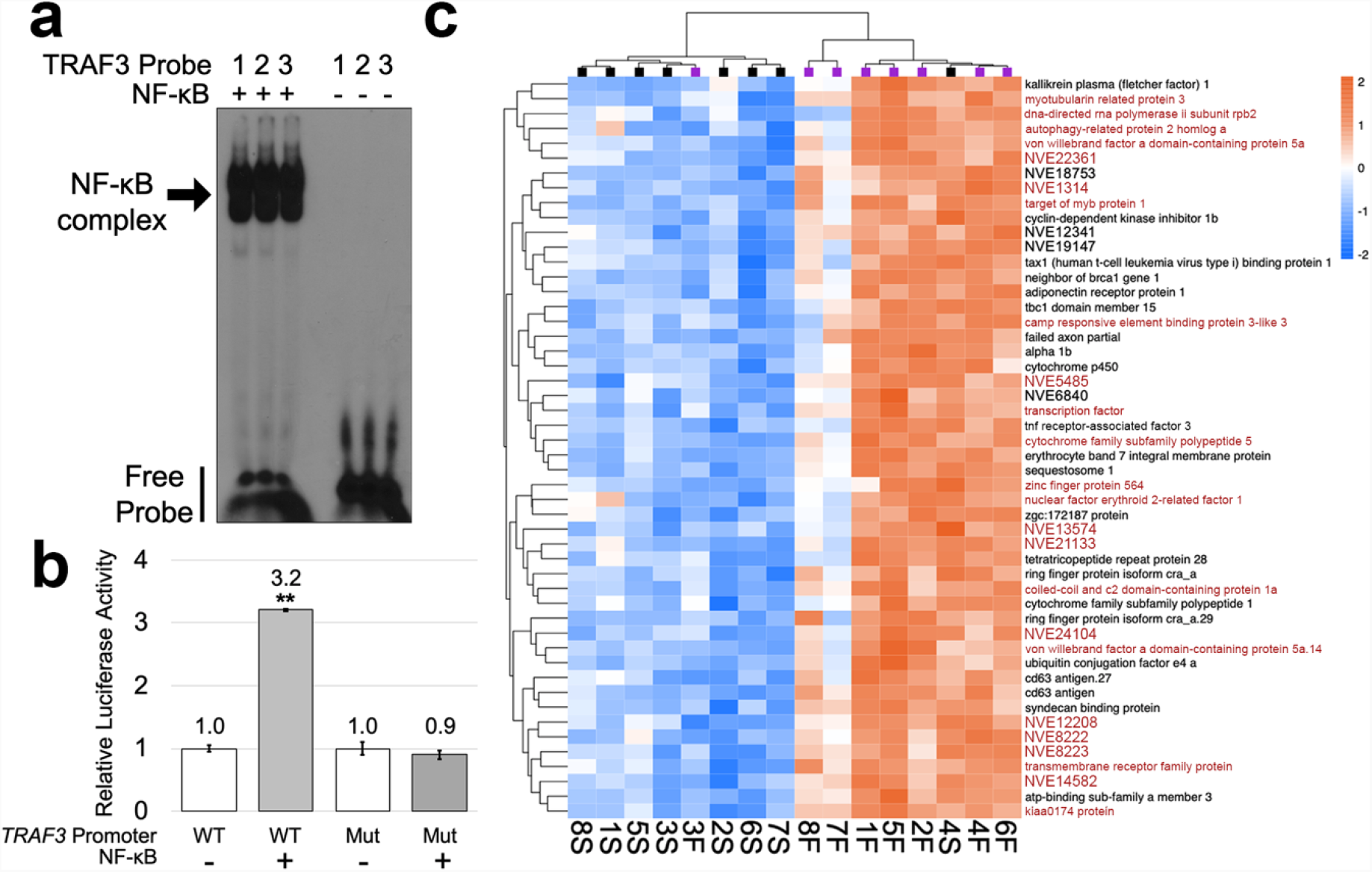
Nv-NF-κB gene expression network analysis. **a**, Nv-NF-κB is able to bind three predicted κB-binding sites (1, 2, 3; see Extended Data Fig. S5) upstream of the *TRAF3* TSS. Bacterially expressed GST-Nv-NF-κB (+) was incubated with ^32^P-labelled probes containing each predicted κB-binding site from the *TRAF3* promoter and analyzed by EMSA. Labelled probes alone (-) were used as negative controls. **b**, Nv-NF-κB can activate an *Nv-TRAF3* promoter luciferase reporter. HEK 293 cells were co-transfected with either an expression plasmid for Nv-NF-κB or the empty vector and luciferase reporter containing either the wild-type *TRAF3* promoter (*TRAF3*-WT) or a mutant *TRAF3* promoter (*TRAF3*-Mut) in which all three NF-κB sites were mutated. Values are averages of triplicate samples and are presented as values relative to the respective empty vector control ± standard error. (**) indicates statistical significance as determined using the Student’s *t test*. **c**, Hierarchical clustered heatmap of “Green” module’s top 50 genes ranked by membership score (kME). Colors indicate direction and magnitude of relative expression (blue, downregulated; red, upregulated). Colored squares at the top indicate feeding regime (purple, fed; black starved). Genes highlighted in red have one or more predicted Nv-NF-κB-binding sites within 1,200 bp upstream of their TSS (see Extended Data Table S3).

To gain a broader overview of a possible Nv-NF-κB gene expression network that is relevant to the starvation response, we used a systems biology-based approach wherein we identified modules of genes whose expression profiles were correlated with NF-κB expression. For this analysis, we used the R package *WGCNA* (Weighted Gene Correlation Network Analysis). This approach generated module eigengenes, or representative gene expression profiles for each module, and we identified the module containing NF-κB (“Green” module). This “Green” module, which contained 1317 genes, had a correlation coefficient of -0.66 with starvation (Extended Data Fig. S3), indicating that expression of genes in this module tended to be downregulated by starvation. We next ranked these transcripts by ‘membership score’ (kME), which is a measure of how strongly each gene corelates with the module’s eigengene. Figure 4c shows a heatmap of the 50 genes in the “Green” module with the highest membership scores. These 50 genes (36/50 annotated) included ones encoding a *TRAF3* homolog and autophagy-related protein 2 homolog A, among others.

To further characterize the genes in the “Green” module, we performed a binary analysis of GO enrichment and found enrichment for terms related to Biological Processes including immunity (i.e., *immune response, regulation of immune response GO:0050776, antigen processing and presentation*), cell signaling (i.e., *regulation of IKK/NF-κB signaling GO:0043122, JNK cascade GO:0007254, MAPK cascade GO:0000165*), and cell death (i.e., *cell death GO:0008219, negative regulation of cell death GO:0060548, regulation of necrotic cell death GO:0010939*) (Extended Data Fig. 4). Additionally, we found that 25 of these top 50 genes had predicted NF-κB binding sites within 1,200 bp of their TSS (Fig. 4c, Extended Data Table S3). Of note, these predicted Nv-NF-κB binding sites have high affinity DNA binding, based on their high Z-scores (Extended Data Table S3) from our previous analysis of DNA binding by Nv-NF-κB using protein-binding microarrays^27^.

## Discussion

Here, we demonstrate a correlation between decreased nutritional status and decreased immunity in the sea anemone *Nematostella vectensis* (*Nv*). We have found that feeding status has a significant impact on gene expression, in addition to the effect of genetic background, consistent with what is seen in other cnidarian gene expression studies involving clonal populations^37,38^. Furthermore, we have shown that the levels and activity of immunity-related transcription factor NF-κB are also reduced under starvation conditions. Thus, we demonstrate a link between nutritional status and immunity in a cnidarian, suggesting that a nutrition-immunity axis has a long evolutionary history. These results also have implications for other cnidarians, e.g., corals, which are endangered by rapidly changing environmental conditions.

Gene expression data and GO enrichment analysis provide insight into the transcriptional effects of starvation in our cnidarian model. Downregulation of terms such as *response to nutrient, lipid metabolic process, carbohydrate metabolic process, glycosylation GO:0070085*, and *proteolysis GO:0006508* under starvation conditions suggest an exhaustion of energetic sources. Downregulation of genes associated with metabolism of different energy sources (carbohydrates, lipids, proteins) during periods of starvation has been previously observed in other species^4,39-42^. These downregulated GO terms indicate that the anemones in our experiments were indeed starved. Moreover, downregulated terms including *ATP biosynthetic process GO:0006754, proton transmembrane transport*, and *electron transport chain GO:0022900* also support the energetic shortage experienced by *Nv* under starvation. Generally, animals that encounter prolonged periods of food deprivation exhibit low metabolic rates^1^, and so it is perhaps not surprising that our data showed downregulation of metabolic processes under food limitation.

We also noted a significant down-regulation of genes and GO terms associated with immunity during starvation of adult *Nv*. Interestingly, food limitation has been previously observed to reduce immunity against bacteria in the caterpillar *Manduca sexta*, as well as decreasing its resistance to oxidative stress^3^, which is a pattern consistent with our GO enrichment for starved *Nv*. Overall, starved *Nv* DEGs showed underrepresentation of GO terms associated with immunity (i.e., *immune response, defense response*, and *antigen processing and presentation*), suggesting a diminished pathogen defense and underrepresented GO terms associated with response to stress (i.e., *reactive oxygen species metabolic process, hydrogen peroxide metabolic process*), suggesting a vulnerability to oxidative stress.

Starved *Nv* polyps had a reduced ability to withstand infection by *P. aeruginosa*, which was correlated with an overall decrease in NF-κB protein levels, as judged by both Western blotting and immunostaining. This relationship between starvation and susceptibility to pathogen infection has been observed in invertebrates^3-5,11^, mice^7,9^, and humans^8,10,43^. A similar relationship has also been observed in *Apis mellifera ligustica* (Italian honeybee), wherein dietary supplementation with an essential fatty acid improved their ability to withstand bacterial infection and resulted in transcriptional upregulation of the NF-κB pathway genes Toll, Myd88, and Dorsal (NF-κB homolog)^11^. The underlying concept shared by all of these examples, as well as our results with *Nv* herein, is that the immune response is an energetically demanding process, which has led to the evolution of proper resource allocation under different nutritional states. For example, *Drosophila* diverts energy from growth and nutrient storage when Toll signaling is activated^44^, and parasitic infection in *Bombus terrestris* (Bumblebee) becomes more virulent under low-nutrient conditions^45^.

Many tropical reef building corals derive the majority of their energy from intracellular symbiotic algae^46^; however, this symbiotic relationship can be lost under a variety of stressors in a process known as coral bleaching^47,48^. Evidence suggests that feeding via heterotrophy is important for corals to mitigate bleaching in the face of warming oceans^17,18^. For example, the branching coral *Montipora capitata* shows enhanced recovery from bleaching compared to other corals by increasing the amount of carbon it acquires through heterotrophy^19^. Similarly, the coral *Pocillopora meandrina* incorporates more heterotrophic carbon when there is more food locally available^49^. Combined with the results presented herein, cnidarians appear to have complex and dynamic ways to respond to stress in the midst of poor nutrient availability.

In addition to bleaching, coral infectious diseases appear to be increasing, which could be due to environmental effects on immunity. Over the last 50 years, ∼40 different coral diseases have been described^50^, with the latest source of concern being Stony Coral Tissue Loss Disease (STCLD) that is affecting Caribbean corals^51,52^. Previous work by our group showed that symbiosis with Symbiodiniaceae algae in the anemone *Exaptasia pallida* has a negative correlation with anemone NF-κB levels, suggesting that the symbiotic state decreases its NF-κB-dependent immunity^27^. Thus, the survival of some cnidarians under certain environmental stressors may be linked to nutrition and immunity.

The phylum Cnidaria emerged approximately 700 million years ago^53^. Since then, individual cnidarians have likely evolved unique genes as part of their immune systems. By taking a systems biology approach, we were able to identify modules of genes that were highly correlated with starvation status in *Nv*. We also identified a module of genes to which NF-κB belongs that was composed of 1317 genes, of which ∼30% are unannotated. Given that many genes in this module are associated with immunity, cell signaling, and cell death, it is likely that some unannotated genes within this same module play roles in immunity, such as being direct anti-microbial effectors. Moreover, we found that 25 of the top 50 genes in the same module as NF-κB had one or more predicted NF-κB-binding sites identified using a PBM-based Nv-NF-κB motif^27^. These results provide avenues to explore novel basal immune gene interactions and are consistent with an evolutionarily conserved role of NF-κB in immunity-related gene regulation.

Previous work in *Nv* identified an unannotated anti-microbial gene that is upregulated in response to the immune stimulatory molecule 2’3’-cGAMP^25^, however, that gene was not affected by starvation-induced downregulation of NF-κB, suggesting that it is not a direct NF-κB target gene. In contrast, several lines of evidence suggest that *TRAF3* is a direct target of Nv-NF-κB: 1) there are three strong NF-κB binding sites located within 1000 bp of the *TRAF3* TSS (Extended Data Fig. 4; Fig. 4a), 2) Nv-NF-κB can activate a reporter locus containing the upstream promoter region of *TRAF3* and this activation requires the upstream NF-κB binding sites (Fig. 4b), and 3) Nv-*TRAF3* is upregulated by activation of the c-GAS-STING pathway in *Nematostella*^25^. It is of interest that mammalian *TRAF3* is a regulator of NF-κB and has a broad role in B-cell immune activation and survival^54^. A correlation between NF-κB and TRAF3 has been reported in several other cnidarian studies. First, NF-κB and *TRAF3* are coordinately induced to high levels in the stony elkhorn coral *Acropora palmata* following acute heat stress^55^; and in *Nv* following 2’3’-cGAMP stimulation of the c-Gas-STING pathway^25^. Additionally, *TRAF3* has been suggested to play a role in coral heat stress response^56,57^, and has been proposed to be a NF-κB target gene in heat-stressed *E. pallida*^58^. Therefore, our results provide a dataset to explore new gene network interactions, as well as leading to the identification of unannotated gene transcripts that are involved in the cnidarian immune system, some of which may be previously unknown anti-microbial agents.

Overall, we show a link between nutrition and immunity in *Nv*, and that NF-κB may play a role in this relationship. These data provide a model for better understanding the interplay between nutrition and certain diseases in cnidarians. The continued study of these important pathways in basal metazoans will further our understanding of where and how these pathways originated, as well as implications for the physiological effects in critical marine organisms as we move into an era of changing climate.

## Materials and Methods

### Care, husbandry and cloning of *Nematostella vectensis*

*N. vectensis* from a Maryland population were obtained from Mark Martindale and Matt Gibson, and spawnings were performed as previously described^12,23,59,60^. Adults, polyps, and larvae were maintained in 1/3 strength artificial sea water (1/3 ASW: ∼12 parts per 1,000) in a dark incubator at 19°C. Adult anemones were fed freshly-hatched brine shrimp (*Artemia)* and young polyps were fed ground *Artemia* in 1/3 ASW three times per week. Water changes were performed weekly for all anemones. To generate clonal pairs, adult animals were allowed to fully relax and were then bisected perpendicularly. Halves were placed into separate wells, and anemones were allowed to regenerate for 30 days. Feeding was paused for all animals during the 30-day regeneration period, i.e., to allow the tentacles to form from the aboral end. Thereafter, both members of the clonal pair were fed in equal amounts.

### RNA extraction and preparation for TagSeq on fed vs. starved anemones

Clonal pairs of adult anemones were generated by bisection and the halves were allowed to heal for 30 days as described above. Thirty days was chosen to allow injury and regeneration-related genes to return to basal levels, as demonstrated previously^13,14^. Clonal pairs were fed equal amounts of food on a regular schedule during healing once all previously aboral ends had developed tentacles. Clonal pairs were split to be fed or starved for 30 days before being flash-frozen on dry ice prior to RNA extraction. Total RNA was isolated from eight clonal pairs with RNAqueous™ Total RNA Isolation Kit (Invitrogen) according to the manufacturer’s instructions, with additional grinding using a plastic pestle during tissue lysis. Next, DNA was eliminated using DNA-*free*™ DNA Removal Kit (Invitrogen). RNA quality was assessed by agarose gel electrophoresis, checking for the presence of ribosomal RNA bands. RNA concentrations were quantified using a NanoDrop ND-1000 Spectrophotometer. Samples were then normalized to 728 ng of total RNA for submission to the University of Texas at Austin – GSAF’s TagSeq Service. Libraries were created by the GSAF and sequenced on a NovaSeq 6000 SR100.

### Transcriptome read mapping

Reads were processed following the TagSeq pipeline (https://github.com/z0on/tag-based_RNAseq). In brief, adapters and poly(A)^+^ tails were trimmed using *Fastx_toolkit* and sequences <20 bp with <90% of bases having quality cut-off scores >20 were trimmed. PCR duplicates sharing degenerate headers were also removed. Resulting quality-controlled reads were aligned to the *Nematostella* transcriptome^60^ using *Bowtie2*.*2*.*0*^62^.

### Differential gene expression and gene ontology analyses

Differential gene expression analysis was performed using DESeq2 v.1.30.1^63^ in R v.4.0.4^64^. The *arrayQualityMetrics*^65^ package tested for outliers, which were defined as any sample failing two or more outlier tests; no outliers were identified. Significant DEGs were identified as those with an FDR-*adjusted p-value* < 0.1. Expression data were normalized using the *rlog* function within the package *vegan*^66^, and normalized data were then used for principal component analysis (PCA) to characterize differences in gene expression between starved and fed (control) groups. Significance was tested by PERMANOVA using the *adonis* function as part of the *vegan* package^66^.

Gene Ontology (GO) enrichment analysis was performed using Mann-Whitney U tests (GO-MWU) based on ranked p-values^67^. GO enrichment results based on the ‘molecular function’ overarching division were plotted as dendrograms with GO categories clustering based on shared genes. Fonts and colors were used to indicate significance and direction of change respectively. To generate a heatmap of “Immune Response”-annotated genes, we used the package *pheatmap*^68^ to showcase differences in expression relative to mean expression across samples.

### Bacterial challenge of *N. vectensis*

Approximately two-week-old polyps were placed into single wells of a 24-well plate, and they were then either fed ground-up *Artemia* for 30 days or starved until infection was initiated. Infection was performed essentially as described previously^25^. Single colonies of *P. aeruginosa* strain PA14 were cultured overnight in Luria Broth, bacteria were centrifuged for 10 min at 1627 x g, rinsed once with 1/3 ASW, centrifuged again, combined, and resuspended to an OD_600_ of ∼0.1 in 1/3 ASW. A small aliquot was taken for plating to calculate CFU/ml. Polyps were infected by placing them in the well of a 24-well plate containing PA14 (1 ml). Survival was monitored daily, mortality was determined based on tissue degradation and the absence of response to light and touch cues^24^. Infection was performed three separate times; once with 12 anemones per feeding regime, and twice with 24 anemones per treatment regime.

### Tissue lysis of *N. vectensis*

Whole protein lysates from anemones were generated following a protocol described previously^69^. Briefly, adult anemones (about 2-cm long) were homogenized using a plastic pestle in 1.5-ml microcentrifuge tubes containing 150 μl of ice-cold AT Lysis buffer with proteinase inhibitors (HEPES [20 mM, pH 7.9], EDTA [1 mM], EGTA [1 mM], glycerol [20% w/v], Triton X-100 [1% w/v], NaF [20 mM], Na_4_P_2_O_7_·10H_2_O [1 mM], dithiothreitol [1 mM], phenylmethylsulfonyl fluoride [1 mM], leupeptin [1 μg/ml], pepstatin A [1 μg/ml], aprotinin [10 μg/ml]). Cell lysis was enhanced by sonicating 5 times for 10 sec on setting 3 with 1 min on ice in between, samples were then passed five times through a 30-gauge needle. NaCl was added to a final concentration of 150 mM. Finally, samples were centrifuged at 13,000 rpm for 30 min at 4°C, and the supernatant was stored at -80°C until needed. For protein lysates from juvenile polyps, 100 40-day old polyps were pooled into a centrifuge tube. Sea water was removed through aspiration and 50 μl 4x SDS sample buffer (Tris-HCl [0.25 M, pH 6.8], SDS [2.3% w/v], glycerol [10% w/v], β-mercaptoethanol [5% v/v], bromophenol blue [0.1% w/v]) was added to the tube. Samples were heated at 95°C for 10 min with vortexing halfway through and at the end. Finally, 50 μl distilled H_2_O was added to the samples.

### Western blotting and electrophoretic mobility shift assay (EMSA)

Western blotting for Nv-NF-κB was performed as previously described^22,26,27,69^. Briefly, proteins were separated on a 7.5% SDS-polyacrylamide gel. Proteins were then transfered at 4°C to a nitrocelluolose membrane at 250 mA for 4 h and then 170 mA overnight. Nitrocellulose membranes were incubated in blocking buffer TBST (Tris-HCl [10 mM, pH 7.4], NaCl [150 mM], Tween 20 [0.1% v/v]) with powdered milk (5% w/v) (Carnation) at room temperature for 1 h. Membranes were then incubated in anti-Nv-NF-κB antibody^22^ (diluted 1:10,000 in blocking buffer) overnight at 4°C. Membranes were washed several times with TBST before incubating with a horseradish peroxidase-conjugated anti-rabbit secondary antiserum (1:4,000, Cell Signaling) for 1 h at room temperature. Membranes were then treated with SuperSignal West Dura Extended Duration Substrate (Pierce), and blots were imaged on a Sapphire Biomolecular Imager. The same filters were also stained with Ponceau S stain to ensure approximately equal total protein loading.

EMSA was performed as previously described^22,26,27,69^ using a ^32^P-labeled κB-site DNA probe (GGGAATTCCC) and adult anemone tissue lysates (described above). Lysates and the κB-site probe were incubated in binding buffer (HEPES [10 mM, pH 7.8], KCl [50 mM], DTT [1 mM], EDTA [1 mM], glycerol [4% w/v]) with poly dI/dC (40 ng) and BSA (200 ng) at 30°C for 30 min. Supershifts were performed by incubating samples with 2 μl of anti-Nv-NF-κB antiserum, after binding to the DNA probe, for 1 h on ice. EMSA gels were dried and then imaged on a Sapphire Biomolecular Imager.

The EMSA for *TRAF3* promoter region was performed as above, except using bacterially expressed GST-Nv-NF-κB, which was purified as described previously^27^. Purified GST-Nv-NF-κB was then incubated in binding buffer as described above with each of the following ^32^P-labeled probes:

(1) 5’-TCGAGAGGTC<u>GGGAAAGCCCC</u>CCCCCG-3’

(2) 5’-TCGAGAGGTC<u>GGGAAACCCCC</u>CCCCCG-3’

(3) 5’-TCGAGAGGTC<u>GGGGAACTCCC</u>CCCCCG-3’

Underlined sequences are predicted NF-κB binding sites in the *Nv TRAF3* promoter (Extended Data Fig. 5). The dried EMSA gel was exposed overnight to X-ray film at -80°C overnight. Film was developed using a standard X-ray film developer.

### Immunohistochemistry of *N. vectensis* polyps

Immunohistochemistry was performed as previously described^22,35,69^. Polyps were fixed in formaldehyde (4%) in 1/3 ASW overnight at 4°C and washed three times with PTx (Triton X-100 [0.2% v/v] in PBS). Antigen retrieval was done by microwaving samples in warm urea (5% w/v) at the lowest setting for 5 min. Samples were cooled at room temperature for 20 min. Samples were then washed three times with PTx. Samples were moved to blocking buffer (PTx + normal goat serum [5% v/v] + BSA [1% w/v]) and allowed to permeabilize overnight at 4°C on a nutator. Blocking buffer was replaced with anti-Nv-NF-κB primary antiserum diluted in blocking buffer (1:100) and incubated overnight at 4°C. Samples were then washed four times with PTx and incubated in Texas-red-conjugated anti-rabbit secondary antiserum (1:160, Invitrogen). Polyps were then washed four times with PTx. Nuclei were stained by adding DAPI to a final concentration of 5 mg/ml. Samples were imaged on a Nikon C2+ Si confocal microscope. NF-κB-positive cells were counted using the *Cell Counter* plug-in in ImageJ.

### Luciferase reporter gene assay

Luciferase reporter gene assays were performed in HEK 293 cells as previously described^22^. Cells were plated in 6-well 35-mm plates to 60% confluence and transfected with: 0.5 μg of (i) pGL3-*Nv*-*TRAF3*, which consists of pGL3 with 1,220 bp of the promoter region of *Nv*-*TRAF3* cloned upstream of the luciferase gene; or (ii) pGL3-*Nv*-*TRAF3*-3X-mut, similar to (i) but with all three putative κB-binding sites mutated to 5’-”GGGGAAAGCTT”-3’, and 2 μg of (i) a Nv-NF-κB expression plasmid or (ii) a pcDNA empty vector. Every transfection was performed with 15 μg of PEI. Two days after the transfection, cells were lysed with Reporter Lysis Buffer (Promega) following manufacturer’s instructions.

### Weighted correlation network analysis (WGCNA)

Weighted Gene Correlation Network Analysis was performed using *WGCNA*^70^ and genes with low basemean values (<3) were removed and all remaining data were *rlog-*normalized. Outlier samples were checked within the *WGCNA* package, and no outliers were detected. Unsigned connectivity between genes was determined and eigengene expression of these modules were correlated to feeding conditions. The “Green” module was chosen by manually searching modules for the NF-κB transcript. Gene Ontology (GO) enrichment analysis was performed as described in the ‘Differential Gene Expression Analysis’ text with the modification of using continuous kME values instead of -log p-values. To generate module heatmap, genes with the highest module membership scores (kME values) within specific modules (e.g., “Green” module) were identified and relative expression was plotted using the package *pheatmap*^68^.

To identify genes with putative NF-κB binding sites, the transcripts of the top 50 “Green genes” (Fig. 4c) were aligned to the *N. vectensis* genome^71,72^ on Ensembl^73^ using BLAST to identify genomic location. We then extracted 1,200 bp upstream of the TSS of every matched gene. To identify Nv-NF-κB binding sites in these upstream regions, we used the program FIMO^74^ (with a p-value cutoff of 1E-04) and a DNA site motif based on Nv-NF-κB DNA binding from a Protein Binding Microarray (PBM) ^27^.

## Supporting information

List of Differentially Expressed Genes

## Acknowledgments

This research was supported by National Science Foundation grants IOS-1354935 and IOS-1937650 (to T.D.G. and S.W.D.). P.J.A.C. was supported in part by a Warren McLeod Marine Fellowship, and N.D. was supported by the Boston University Undergraduate Research Opportunities Program. We thank Mark Martindale (Whitney Laboratory for Marine Science) and Matt Gibson (Stowers Institute for Medical Research) for sharing *Nv* with us, Stephen Lory (Harvard University) for the PA14 strain, and Trevor Siggers (Boston University) for help with the analysis of Nv-NF-κB DNA-binding sites.

## Competing interest statement

The authors declare no competing interest.

## Author contributions

P.J.A.C. performed immunohistochemistry, infection experiments, Western blotting and EMSA of *Nv* lysates. P.J.A.C. and N.D. performed Western blots and EMSAs. P.J.A.C., J.F., and S.W.D. contributed to writing scripts used in this manuscript. J.J.B. performed luciferase assays and *TRAF3* EMSA. P.J.A.C., S.W.D., and T.D.G. designed experiments and wrote the manuscript.

## Notes

### Competing Interest Statement

The authors have declared no competing interest.

