## Supplementary Information for "Starvation Decreases Immunity and Immune Regulatory Factor NF-κB in the Starlet Sea Anemone *Nematostella vectensis*"

Thomas D. Gilmore

**This file includes:**

Figures 1 to 5

Tables 1 to 3

**Other materials for this manuscript include the following:**

Dataset 1

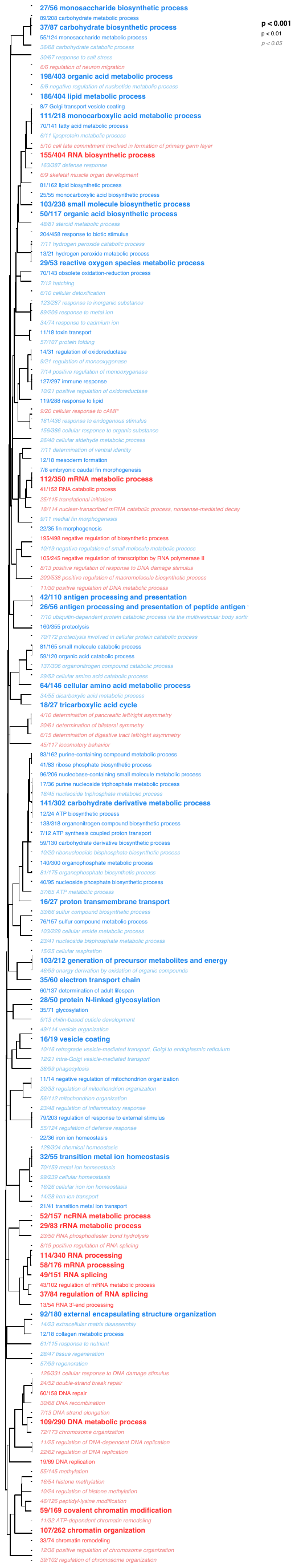

Fig. 1. Gene Ontology (GO) analysis of significantly enriched ‘Biological Process’ terms in starved anemones using Mann-Whitney U tests (GO-MWU) based on ranked p-values. Dendrograms clusters terms based on genes shared between categories. Direction of enrichment is indicated by color; blue indicates underrepresented terms, and red indicates overrepresented terms in starved anemones. Font indicates p-values as indicated at the top right.

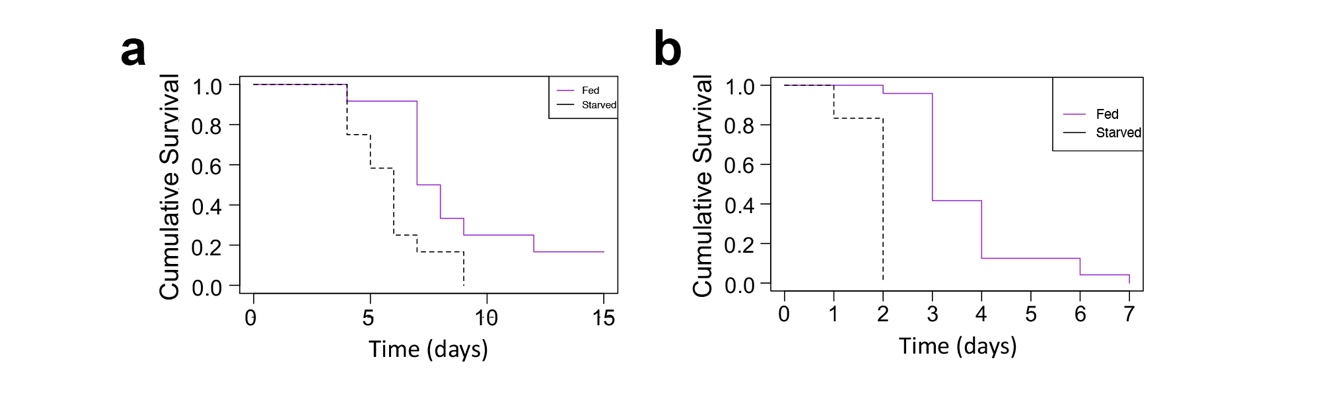

Fig. 2. Starved anemones have increased susceptibility to *Pseudomas aeruginosa* infection-induced death. Two-week old anemones were either fed on a regular schedule (purple) for 30 days or starved (black), and were then infected with a, 4.4 x 10^8^ or b, 2.5 x 10^8^ CFU/ml of *P. aeruginosa* at 28˚C. Survival was monitored daily for 15 days and recorded. a, N = 12 and b, N = 24 for each condition. Significance was determined using Kaplan-Meier statistics.

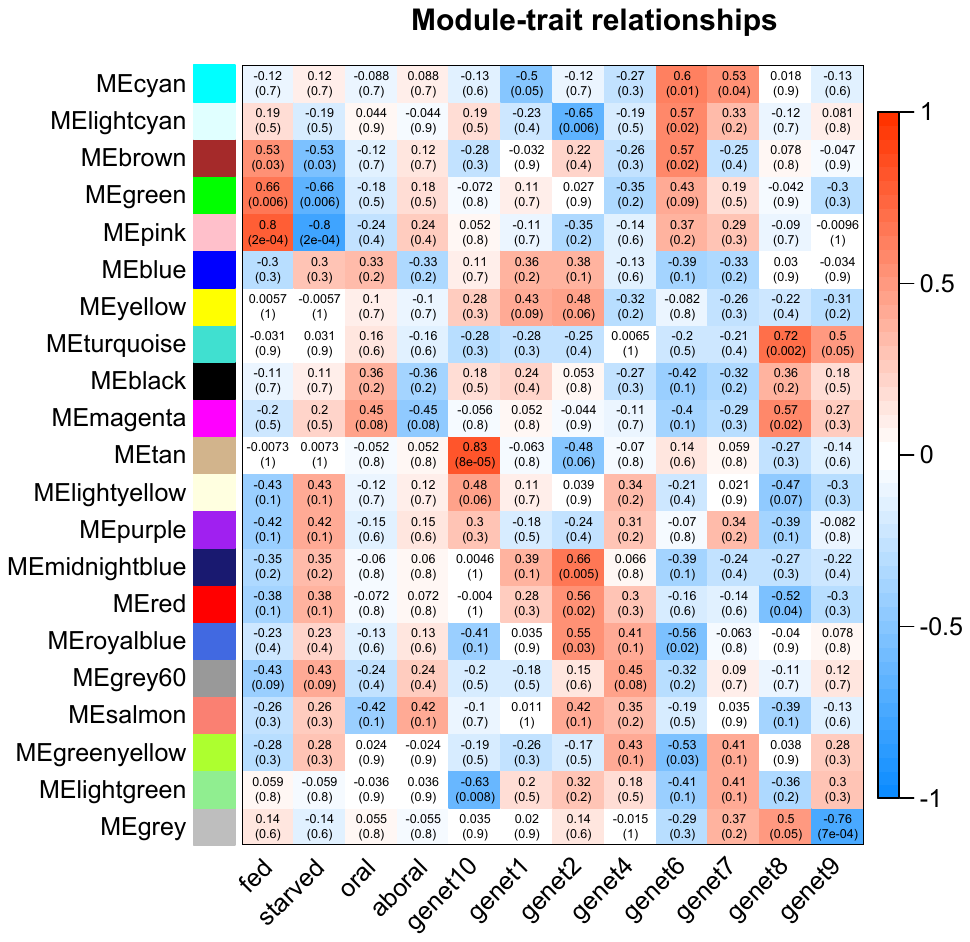

Fig. S3. WGCNA-generated heatmap of traits and gene modules. WGCNA was performed using the package *WGCNA*. Color and shading correspond to correlation between trait (bottom) and gene module (left), positive numbers (red) indicate positive correlation, negative numbers (blue) indicate negative correlation between modules and traits. ‘fed’ and ‘starved’ indicate feeding status; ‘oral’ and ‘aboral’ indicate whether individuals originated from oral or aboral end; genets 1-8 indicate clonal genotype.

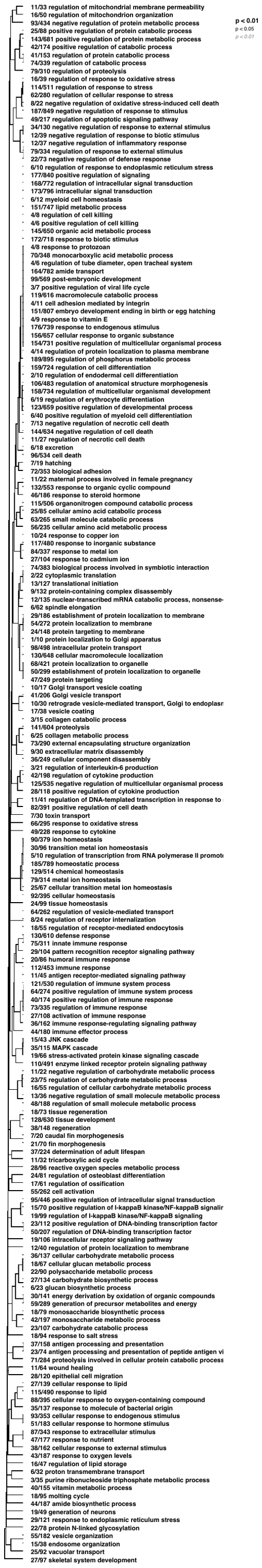

Fig. S4. Gene Ontology (GO) analysis of significantly enriched ‘Biological Process’ terms in WGCNA-generated gene module containing NF-κB using Mann-Whitney U tests (GO-MWU) based on continuous kME. Dendrograms clusters terms based on genes shared between categories.

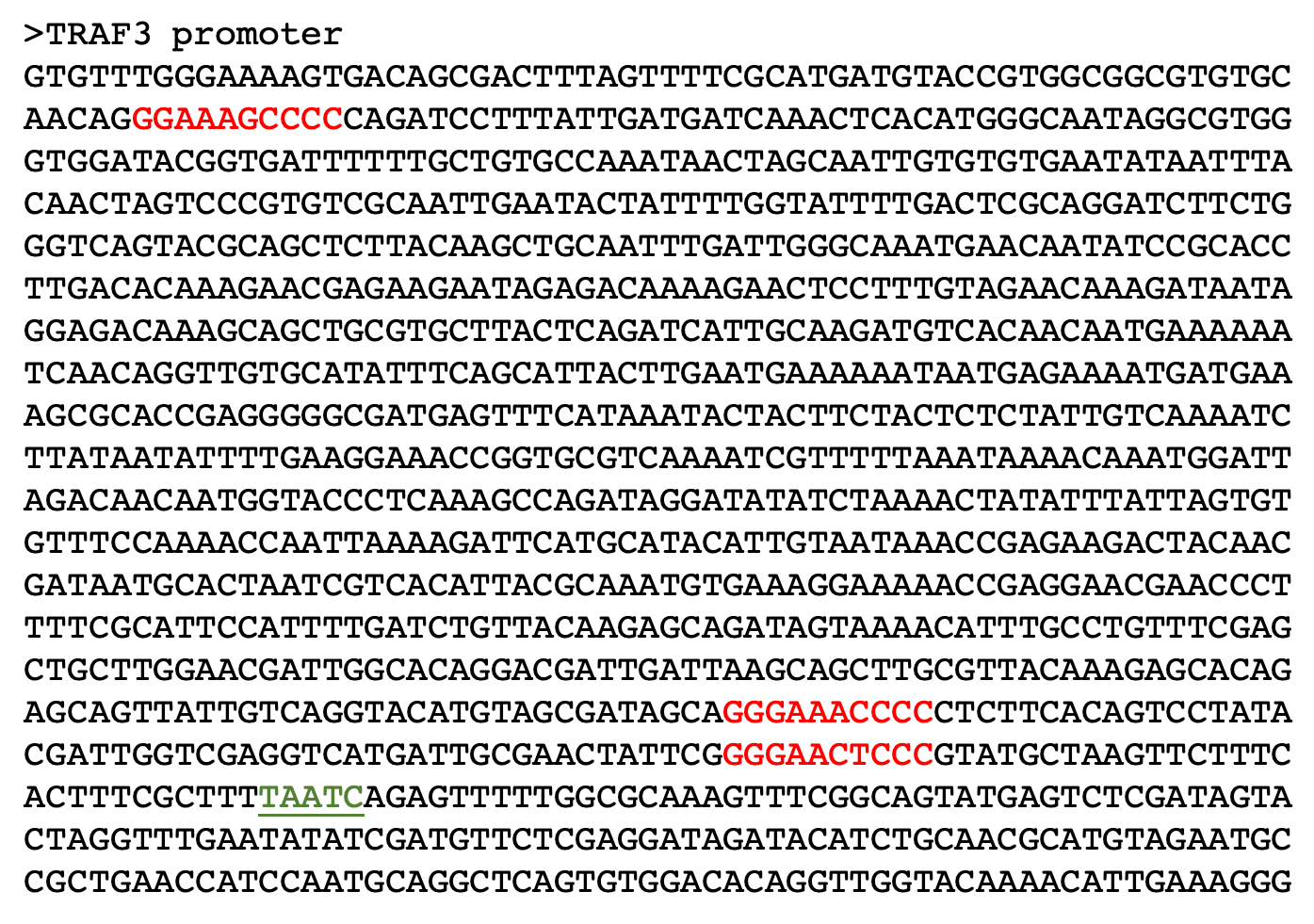

Fig. S5. The *Nv* *TRAF3* proximal promoter region. Predicted NF-κB binding sites are highlighted in red. The three sites are numbered 1, 2, and 3 from top to bottom. These sites had PBM-based DNA-binding Z-scores as follows: site 1, 6.1; site 2, 9.7; and site 3, 6.9 (see ref.^27^ for details). The transcription start site is highlighted in green and underlined.

Table S1. Sample metadata for anemones used in differential gene expression analysis. Table includes sample names, clonal origin, what end (oral/aboral) the anemone regenerated from, feeding regime, file label for sequencing data available on SRA PRJNA837630, and total raw reads for each sample.

| Anemone | Clone parent | Origin end | Feeding regime | File label | Raw reads |
| --- | --- | --- | --- | --- | --- |
| 1F | 1 | Oral | Fed | 1OF | 8,249,165 |
| 1S | 1 | Aboral | Starved | 1AS | 6,043,016 |
| 2F | 2 | Oral | Fed | 2OF | 5,891,586 |
| 2S | 2 | Aboral | Starved | 2AS | 6,095,160 |
| 3F | 3 | Oral | Fed | 4OF | 9,197,316 |
| 3S | 3 | Aboral | Starved | 4AS | 7,487,177 |
| 4F | 4 | Aboral | Fed | 6AF | 5,402,652 |
| 4S | 4 | Oral | Starved | 6OS | 5,448,963 |
| 5F | 5 | Aboral | Fed | 7AF | 5,538,452 |
| 5S | 5 | Oral | Starved | 7OS | 7,403,004 |
| 6F | 6 | Aboral | Fed | 8AF | 5,215,428 |
| 6S | 6 | Oral | Starved | 8OS | 8,528,160 |
| 7F | 7 | Aboral | Fed | 9AF | 7,597,310 |
| 7S | 7 | Oral | Starved | 9OS | 6,244,255 |
| 8F | 8 | Aboral | Fed | 10AF | 6,604,076 |
| 8S | 8 | Oral | Starved | 10OS | 8,152,984 |

Table S2. Whole-mount immunofluorescence was performed on 40-day-old polyps that were either fed for 30 days or never fed. Nv-NF-κB was detected with a primary anti-Nv-NF-κB antiserum and Texas Red-labeled secondary antiserum, and nuclei were detected with DAPI. Samples were imaged on a Nikon C2+ Si confocal microscope. NF-κB-positive cells were counted using the *Cell Counter* plug-in in ImageJ.

| **Feeding Treatment** | **1** | **2** | **3** | **4** | **5** | **6** |
| --- | --- | --- | --- | --- | --- | --- |
| Fed | 213 | 91 | 169 | 169 | 411 | 140 |
| Starved | 111 | 146 | 49 | 91 | 19 | 62 |

Table S3. “Green” module genes with FIMO-predicted Nv-NF-κB-binding sites. The top 50 genes ranked by membership score (kME) (Fig. 4c) where aligned to the *Nv* genome using BLAST, and sequences 1,200 bp upstream of the TSS were extracted. Putative Nv-NF-κB-binding sites were predicted using a PBM-generated binding motif and FIMO with a p-value cutoff of 1E-04. Gene names are provided where available as annotated in the transcriptome^60^, otherwise transcript names are given. PBM Z-score for binding sites from ref. 27 are provided where available. Of note, NVE8222 and NVE8223 aligned to the same region of the genome and are likely different transcripts of the same gene.

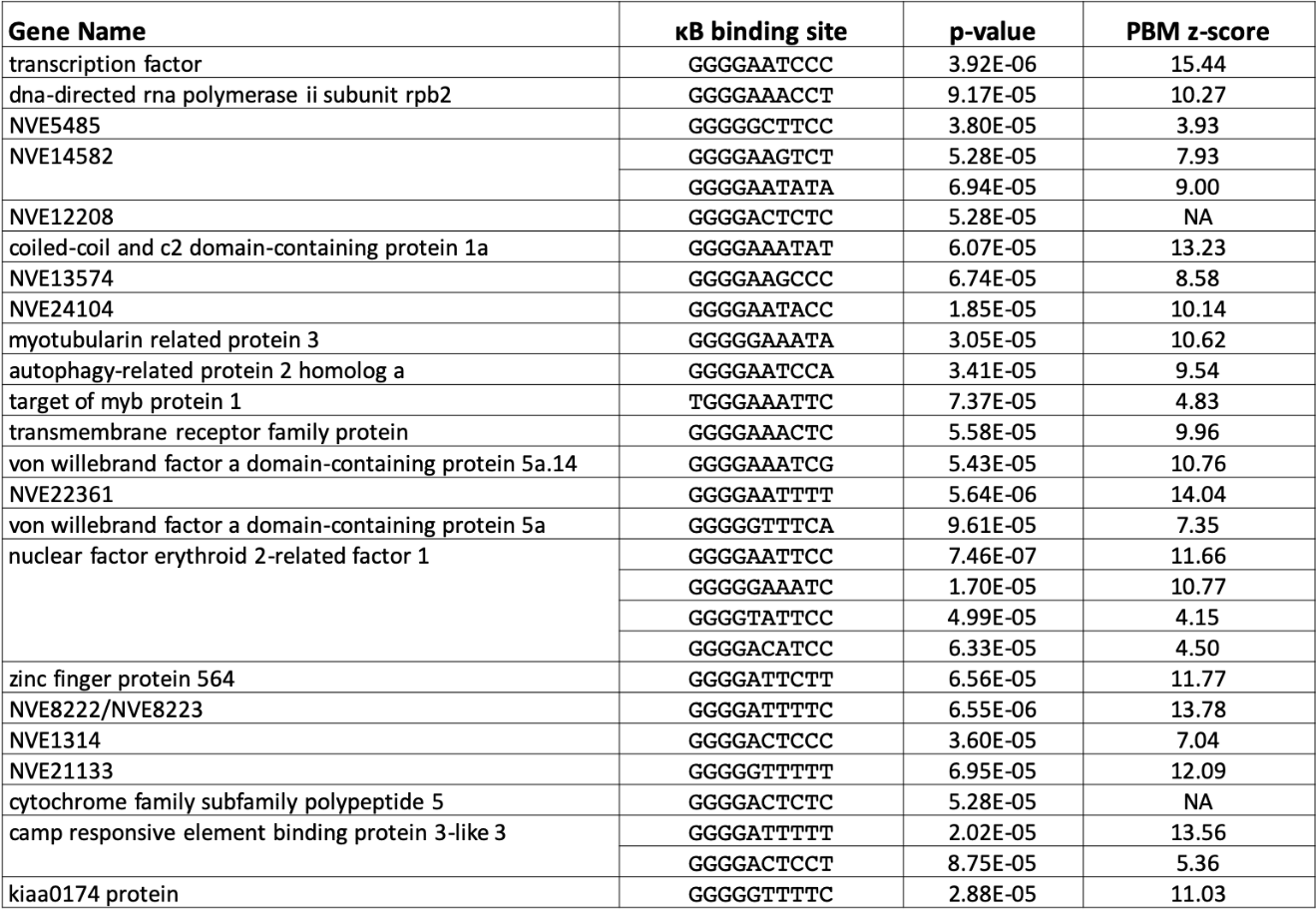

Dataset S1 (separate file). List of significant DEGs (*FDR adjusted p-value* < 0.1) in starved anemones relative to fed controls. Table includes output data generated by *DESeq2*.
